# MOSGA 2: Comparative genomics and validation tools

**DOI:** 10.1101/2021.07.29.454382

**Authors:** Roman Martin, Hagen Dreßler, Georges Hattab, Thomas Hackl, Matthias G. Fischer, Dominik Heider

## Abstract

Due to the highly growing number of available genomic information, the need for accessible and easy-to-use analysis tools is increasing. To facilitate eukaryotic genome annotations, we created MOSGA. In this work, we show how MOSGA 2 is developed by including several advanced analyses for genomic data. Since the genomic data quality greatly impacts the annotation quality, we included multiple tools to validate and ensure high-quality user-submitted genome assemblies. Moreover, thanks to the integration of comparative genomics methods, users can benefit from a broader genomic view by analyzing multiple genomic data sets simultaneously. Further, we demonstrate the new functionalities of MOSGA 2 by different use-cases and practical examples. MOSGA 2 extends the already established application to the quality control of the genomic data and integrates and analyzes multiple genomes in a larger context, e.g., by phylogenetics.

## Introduction

The constantly increasing number of available wholegenome sequences, mainly derived by high-throughput sequencing, provides more and more biological insights (1). In particular, the sequencing of new bacterial, archaeal, and viral genomes is now routine practice. It has accelerated discoveries in microbial diversity and evolution, providing new insight into microbiome function, human health, and biogeochemical cycling (2–6). However, the generation of high-quality eukaryotic genomes has been hampered in the past by sequencing costs and assembly complexity, limiting sequenced eukaryotes to those of medical or economical interest (7). More recently, advances in Illumina high-throughput sequencing and emerging long-read sequencing technologies developed by Pacific Biosciences and Oxford Nanopore have also rendered the generation of genome assemblies of large eukaryotes a routine task (8–10). These proceeding trends in data availability, advanced techniques, and improved affordability require more facilitated access for scientists to analyze these sequence data. To address the challenging issue of correctly annotating new genomes, we developed the Modular Open-Source Genome Annotator (MOSGA) (11). While the number of available genomic data increases continuously, the quality of these data is as important as their availability. Low-quality genome assemblies will introduce errors (12) and, as such, affect the annotation quality. In addition, genome assemblies can be incomplete or contaminated, which does not necessarily affect the annotation process but will hinder the submission process and may lead to wrong conclusions. This study presents the new functionalities in MOSGA 2, such as new workflows for comparative genomics and the introduction of validation tools into the framework. In brief, MOSGA 2 incorporates comparative genomic workflows derived from Hackl et al. (13) and validation tools from Pirovano et al. (14).

We initially developed MOSGA to facilitate genome annotation, including an easy-to-use web interface based on a robust, scalable, modular, and reproducible platform. Besides new genome annotations, genome assemblies or multiple genomes can be placed in a bigger context like comparative genomics. Therefore, we integrated several tools into MOSGA 2 to calculate and output the phylogenetic trees, the average nucleotide identity, the completeness of the genomes, and the protein-coding gene comparison based on previously performed protein-coding gene annotations. In addition, we incorporated tools to identify the occurrence of contamination and present a new scaffold-detection method for organellar sequences present in genome assemblies.

## Software changes

### Phylogenetics

MOSGA 2 integrates a workflow based on nine established tools to preprocess and compute phylogenetic trees on multiple genomes. Preprocessing steps include selecting a database for genes, constructing and trimming multiple sequence alignment (MSA).

Therefore, we included BUSCO (15, 16) and EukCC (17) to identify single-copy genes in genomes for the phylogenetic computation. The user has to choose from one of these tools or even combine both as potential phylogenetic markers. MOSGA 2 concatenates the chosen gene marker sequences into a new FASTA file. The user has to select a program for the MSA construction, such as MAFFT (18, 19), ClustalW (20, 21), or MUSCLE (22). For an optional MSA trimming, we integrated trimAl (23) and ClipKIT (24). Phylogenetic relationships are then reconstructed by maximum likelihood or distance algorithms using RAxML (25) or FastME (26), respectively, and the resulting tree are visualized by ggtree (27) and ape (28). Tree rooting is optional and can be defined by marking an uploaded genome assembly as an outgroup.

### Average nucleotide identity

For whole-genome similarity metrics, MOSGA 2 includes a comparative genomics workflow that calculates the Average Nucleotide Identity (ANI) across all genomes. To achieve this, we integrated FastANI (29), which compares the genomes against each other. Further, the ANI values are represented as a heatmap.

### Protein-coding genes comparison

MOSGA 2 integrates a comparative genomics workflow that compares proteincoding genes from all uploaded genomes against each other. This comparison requires a previous annotation of proteincoding genes, which can be achieved via the annotation pipeline or by importing already annotated genomes as GBFF (GenBank flat format) files. This allows, for example, a comparison between different gene prediction tools or between reference and experimental annotations. Technically, MOSGA 2 extracts the protein-coding sequences and matches them against each other. Matches above a defined threshold will be binned back to a genome, and the average coding content similarities between the genomes are displayed as a heatmap. This analysis allows consistency checks for gene predictions across different genomes. This method compares the nucleotide sequence of protein-coding genes and has similarities with the concept of Average Amino Acids Identity.

### Genome completeness

The identification of single-copy genes is a crucial step for phylogenetic analysis. Therefore, we integrated BUSCO and EukCC into MOSGA 2. While BUSCO’s data source is OrthoDB (31), EukCC relies on PANTHER (32). These tools can estimate the completeness of assemblies, and MOSGA 2 integrates them for validation into the annotation and the comparative genomics workflow. Genome completeness results for each genome are visualized together in the comparative genomics workflow and the annotation workflow separately.

### Contamination detection

To detect potential contamination in a genome assembly, such as sequences from other organisms or residual sequencing adapters, we integrated two validation tools to identify putative contamination. In the case of a gene prediction workflow with RNA-seq based gene prediction, MOSGA 2 offers BlobTools (33) to estimate the taxonomical source of each scaffold. Such sources may be helpful to identify putative biological contaminants. Additionally, MOSGA 2 includes NCBI’s VecScreen based on the UniVec database to identify adapter sequences.

### Organellar DNA scanner

MOSGA 2 is optimized for annotating nuclear DNA sequences from eukaryotic cells and is less suited for organellar DNA, such as mitochondrial and plastid genomes. To identify organellar DNA in genome assemblies, MOSGA 2 combines information from GC-content, plastid and mitochondrial reference protein databases with RNA prediction algorithms such as barrnap and tRNAscan-SE 2.0 (34). At the end of each annotation job, MOSGA 2 creates a relative ranking with the most likely scaffolds. The scoring is an arithmetic calculation based on the density of organellar-specific genes, the numbers of tRNAs, rRNAs, and the GC-content variance. To create the plastid and mitochondria reference proteins databases, we clustered respective RefSeq (35) databases with MM-Seq2 (36) and only kept representative sequences of these clusters into our databases to remove redundancy.

### Taxonomy search

In several MOSGA annotation jobs, we observed that multiple users did not select the best suitable gene-prediction models for the given data. Depending on the selection of gene prediction tools, this task could be challenging since, for example, the gene predictor Augustus (37) currently includes already 80 species-specific models. Identifying the most suitable models requires knowledge or an educated guess about each listed species and their relatedness. To support the user during this task, we implemented a taxonomy search for taxonomy-related options. To do so, users select the species name for the uploaded genome assemblies and MOSGA 2 searches for each tool’s best putative species- or lineage-specific parameter. Internally, MOSGA 2 contains a trimmed version of the NCBI taxonomy database (38) and searches for the shortest weighted distance between two given nodes. This feature is available for the gene-prediction tools Augustus, GlimmerHMM (39), and SNAP (40) and the validation tool BUSCO.

### Annotation quality

By default, each finished MOSGA genome annotation is validated by NCBI’s tbl2asn. In MOSGA 2, we inserted additional multiple filters that improve NCBI compatibility, mainly following Pirovano et al. (14). We integrated additional filters checking the suggested sizes for exons, introns, and the completeness of proteincoding sequences; this includes internal stop-codons and correct start- and stop-codons.

### Integration of existing annotations

MOSGA 2 can import existing genome annotations in GenBank flat format (GBFF) and, therefore, can combine or complete existing annotations with output from additional prediction tools. Results from prediction tools can be visually compared using JBrowse (41). The GBFF file support is not limited to annotation jobs, but can also be used for comparative genomics tasks or to mix different file formats, which are subsequently interpreted by our tool.

### External Application Programming Interfaces

As another new feature, we introduced three APIs to established external tools: g:Profiler g:GOST (42) for functional enrichment analysis, Integrated Interactions Database (43), and the STRING database (44) for Protein-Protein Interactions Analysis. MOSGA 2 submits predicted protein identifiers from functional annotation to these tools by enabling multiple APIs in the annotation mode and passing the results back into the job submission.

### Gene-prediction

To improve one of the main tasks of MOSGA, we included two new workflows to predict proteincoding genes with BRAKER 2 (45). New genes can be found based on protein- or orthology-based evidence derived from the OrthoDB database.

## Results

To demonstrate the usability of MOSGA 2, we will demonstrate several features with exemplary cases.

### Phylogenetics, genome completeness and Average Nucleotide Identity

We performed a phylogenetic analysis based on seven different genome assemblies from the *Saccharomyces* genus (see Table S1). BUSCO served as the source for gene detection with the Eukaryota lineage OrthoDB dataset. MOSGA 2 concatenated the genes, produced a multiple-sequence alignment with MAFFT, and trimmed the MSA with trimAl. The trimmed MSA is used to calculate the phylogenetics with RAxML. The resulting phylogenetic tree in Figure 1 displays a branching topology that is identical to previous analyses (30). We marked *S. uvarum* as an outgroup to define the tree rooting. Since BUSCO provided the gene source for the phylogenetic analysis in this example, MOSGA 2 evaluated the genome completeness for each genome that is shown in Figure 2. The distribution of common missed and unique missed BUSCOs for each genome is shown in Figure S2. Most of the missing BUSCOs were common to all genomes, indicating that the Eukaryota lineages do not entirely cover all species from the *Saccharomyces* genus or that these orthologs are indeed absent from *Saccharomyces* genomes. According to the BUSCO genome completeness analysis shown in Figure 2, the genome assembly of *S. mikatae* seems to be incomplete. This could be confirmed by applying an additional EukCC analysis for genome completeness (see Figure S1). The incompleteness of *S. mikatae* is likely to be derived from the high number of scaffolds and indicates a low-quality genome assembly. In this context, EukCC detected a maximum silent contamination value over 40 % for this assembly. Furthermore, MOSGA 2 calculated the Average Nucleotide Identity that is shown in Figure 2. *S. cerevisiae’s* genome shows a high similarity with to *S. paradoxus* genome and *S. eubayanus* to *S. uvarum.* Indeed, phylogenetic tree analysis indicates a closer relatedness of these species.

**Fig. 1.**
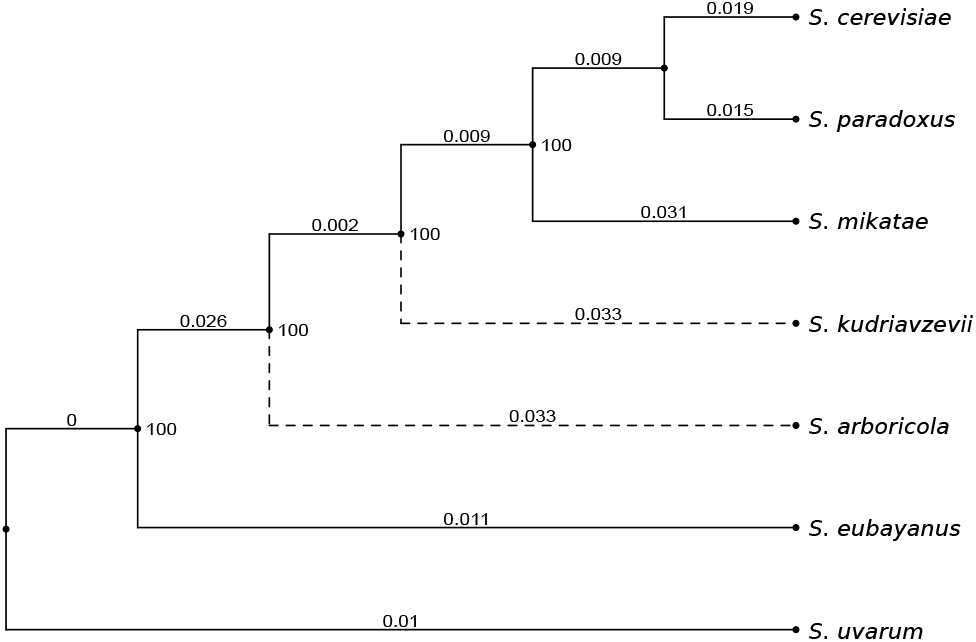
Multi-gene phylogenetic tree of seven *Saccharomyces* species. This phylogenetic tree output of MOSGA 2 was computed using BUSCO MAFFT, traimAl, and RAxML based on 173 identified BUSCO genes. The tree topologies are identical to the one described in Peter et al. (30). Dashed lines indicate a higher distance than the 90th quantile.

**Fig. 2.**
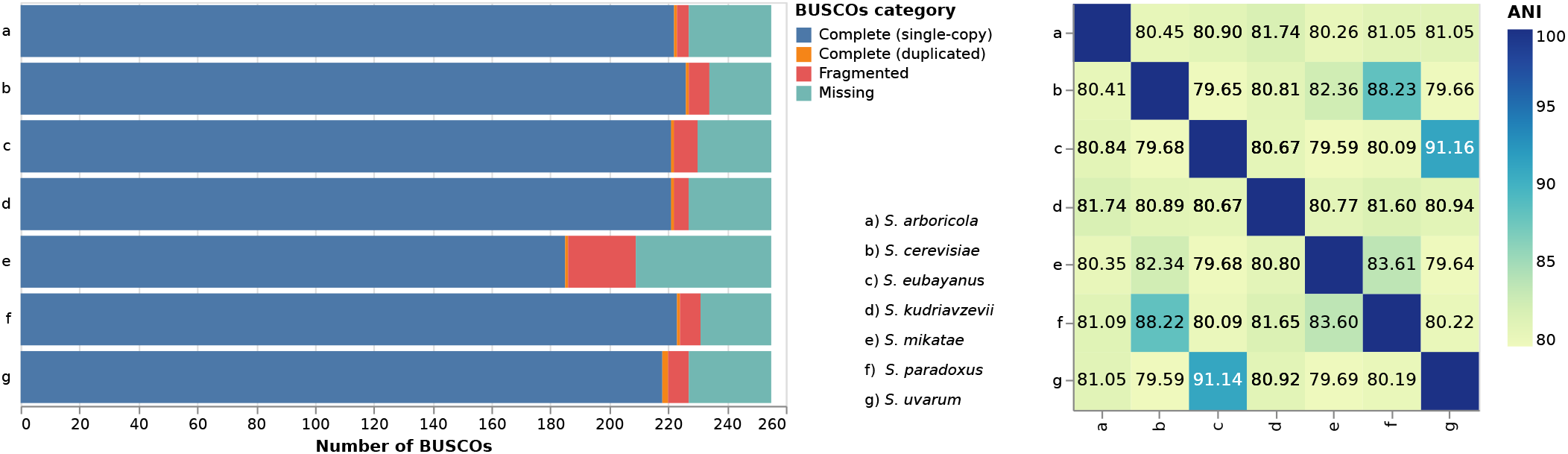
Merged visualization of BUSCO and FastANI results. Visual representation from the BUSCO analysis based on the Eukaryota OrthoDB lineage shows the genomecompleteness of seven different yeast species and the heatmap from Average Nucleotide Identity (ANI) analysis.

### Protein-coding genes comparison

To demonstrate the protein-coding gene comparison feature of MOSGA 2, we used Augustus v3.4.0 to predict protein-coding sequences in *S. paradoxus* and five *S. cerevisiae* strains. The chosen assemblies are listed in Table S2. Protein-coding prediction consistency can be checked independently from the used software, especially for annotating different strains from the same species or identifying contaminant species. Depending on the selected threshold for acceptance of a gene match, the strictness increases, and the genes for each genome are mostly only matching with their original genome, as shown in Figure 3. By decreasing the identity threshold, the total matches became more undifferentiable. Yet, it is important to note that it is possible to differentiate between *S. paradoxus* and *S. cerevisiae* genes similarities. Additionally, we could observe that *S. cerevisiae* strain SK1 has more gene similarities with S288C, while HLJ167, Y12 and sake001 are more similar to each other. We could confirm this observation by calculating the phylogenetic tree with 1873 BUSCOs for these genomes based on the BUSCO *Saccharomycetes* OrthoDB data source, shown in Figure S3.

**Fig. 3.**
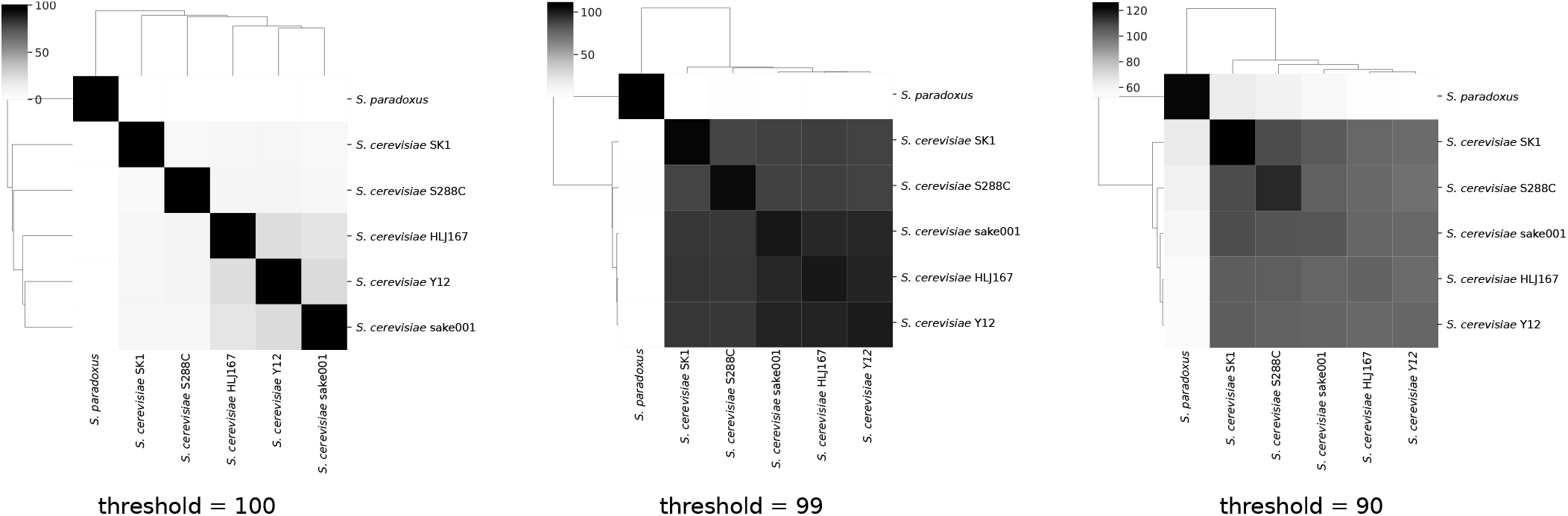
Gene similarity matrix of six different *Saccharomyces* strains and species based on Augustus predicted protein-coding genes. Depending on the selected threshold for the binning, the similarities differentiate more or less strongly between each genome. Values are relative and normalized by the genome sizes.

### Organellar DNA scanner

We applied our organellar DNA scanner on twenty diverse eukaryotic genomes to quickly identify the mitochondrial, chloroplastid, or other organelles’ scaffolds. We chose genome assemblies from various eukaryotic taxa, including plants, nematodes, protists, fungi, vertebrates, and mammalians, to evaluate our scoring. For that purpose, we performed for each genome individual annotations jobs with default settings but without any proteincoding gene prediction. The results of the organellar DNA scanner are represented in a table containing all scaffolds and putative indicators for the presence of an organellar scaffold. To facilitate the interpretation of this table, we introduced a simple scoring system that considers all putative indicators. The highest-ranking score sorts the resulting scoring table rows, shown for *Nannochloropsis oceanica* in Table S3. In 16 out of 20 assemblies, the first hit belonged to one of the organellar scaffolds. An overview of the results is shown in Table 1 and a detailed table considering the type of the organelles is represented in Table S4. The scoring generally performs worse if the number of unplaced contigs increases. We validated the positive matches by comparing our identified first-hit scaffold with the NCBI Genome database.

**Table 1.** Summarized results from the organellar DNA scanner of twenty eukaryotic genome assemblies. Asterisks mark assemblies that contain multiple unplaced scaffolds. **+++** identifies an organellar scaffold as the first suggestion, **++** identifies an organellar scaffold within the first one percent of the suggestions, **+** identifies an organellar scaffold within the first five percent of the suggestions. The total result ranking is shown in Table S3.

| Species | No. Scaffolds | Result |
| --- | --- | --- |
| *Arabidopsis thaliana* | 7 | +++ |
| *Apis mellifera* | 11 | +++ |
| *Babesia microti* | 6 | +++ |
| *Bos taurus* | 2211 | +++ |
| *Caenorhabditis elegans* | 7 | +++ |
| *Cafeteria burkhardae* | 170 | +++ |
| *Cardiosporidium cionae* | 2204 | ++ |
| *Corvus cornix* | 113 | +++ |
| *Danio rerio* | 1917 | +++ |
| *Drosophila melanogaster* | 1870 | +++ |
| *Homo sapiens* | 164 | +++ |
| *Ipomoea triloba* | 17 | +++ |
| *Nannochloropsis oceanica* | 32 | +++ |
| *Plasmodium falciparum* | 15 | +++ |
| *Prunus dulcis* | 692 | ++ |
| *Saccharomyces cerevisiae* | 17 | +++ |
| *Salmo salar* | 241573 | ++ |
| *Strongylocentrotus purpuratus* | 871 | +++ |
| *Tribolium castaneum* | 2149 | ++ |
| *Zea mays* | 687 | + |

### Taxonomy search

The integrated taxonomy search in MOSGA 2 tries to identify the most suitable model or lineage for the annotation workflow. Depending on the completeness of the NCBI taxonomy and the available models, it can quickly identify the putative best parameter for each tool. For example, defining *Saccharomyces cerevisiae* as the input genome species, MOSGA 2 selects the *Saccharomyces* Augustus species model and the *Saccharomycetes* OrthoDB data source for BUSCO. As another example, a search for *Taphrina alni* results in an Augustus model match for *Pneumocystis jirovecii* (Figure S4). Both species belong to the subphylum *Taphrinomycotina*. The search depends on the completeness of the NCBI-defined taxonomy. In other cases, if the distance is similar, the selected match will be the first hit.

## Discussion

In MOSGA 2, we streamline the annotation process by providing validation methods and tools. New functionalities enable user-friendly analyses of genome completeness, contamination, or optimal selected parameters. While improving the annotation process, we additionally implemented comparative genomics workflows and thus extended the scope of MOSGA’s capabilities.

We demonstrated that MOSGA 2 generates reliable phylogenetic results using the *Saccharomyces* genus as an example. Depending on the chosen parameters, variances have to be expected, but in this analysis, we only used the default MOSGA 2 settings and selected BUSCO as the gene source. We presented that the protein-coding genes comparison analysis helps to identify differences in samples based on previously predicted protein genes or inconsistent gene predictions.

Furthermore, besides quality controls such as the genome completeness analysis, we showed that MOSGA 2 could facilitate data preparation by identifying organellar scaffolds inside a genome assembly. In most cases, MOSGA 2 could identify the organellar scaffold directly, although this depends on the assembly quality. The precision of the organellar DNA scanner decreases in samples with many scaffolds, which could indicate problems with biological contamination. Moreover, we demonstrated that the taxonomy search feature supports the user in finding the most appropriate model. This search generally relies heavily on NCBI taxonomy quality. This is especially relevant when taxonomical information is incomplete.

During MOSGA 2 development, we implemented feedback and observations from MOSGA users, such as wrongly chosen species models or eukaryotic assemblies with organellar scaffolds. MOSGA 2 improved in terms of quality and user-friendliness by implementing validation tools and functions like the taxonomy search and the organellar DNA scanner. Furthermore, we integrated new workflows that allow comparative genomics analyses and expand the scope of MOSGA analysis for a wider range of applications. Our comparative genomics workflows are based on state-of-the-art tools. For output visualization, we have followed the guidelines of Hattab et al. (46), and for improved genome annotation quality and validation tools, the advice of Pirovano et al. (14). In total, we increased the number of implemented workflow rules from 63 to 129. Figure S5 and Figure S6 show two job examples for comparative genomics and the annotation workflow. Although MOSGA 2 grows in complexity, it relies on our established architecture that allows high scalability and reproducibility thanks to the multiple-layer design and the Snakemake workflow engine (11). We implemented several Snakemake and comprehensive data analysis tests that GitLab performs on source-code contribution to maintain the code quality. We encourage scientists to send us implementation and workflow suggestions to support the development of MOSGA 2.

## Data availability

Our source code is MIT-licensed freely available on GitLab (gitlab.com/mosga/mosga) and Zenodo (doi: 10.5281/zenodo.5121228). An online version of MOSGA 2 is available under mosga.mathematik.uni-marburg.de. This website is free and open to all users, and there are no registration requirements. The calculated results for the given examples are available under mosga.mathematik. uni-marburg.de/phylo for the *Saccharomyces* phylogenetics example, mosga.mathematik.uni-marburg.de/genecomp for the *Saccharomyces* strains gene comparison and mosga. mathematik.uni-marburg.de/organelles for the organellar DNA scanner.

## Competing interests

There is NO Competing Interest.

## Funding

This work was supported by the LOEWE program of the State of Hesse (Germany) in the MOSLA research cluster.

## Author contributions statement

R.M. wrote the manuscript, designed and developed the framework. H.D. implemented the comparative genomics workflows and assisted in the development. G.H. supported the tool development and data visualizations, he proofread and revised the manuscript. T.H. provided the database for the organellar DNA scanner and ideas for implementation and revised the manuscript. M.G.F. revised the manuscript. D.H. supervised the project, discussed the results, and revised the manuscript. All authors read and approved the final manuscript.

## ACKNOWLEDGEMENTS

The authors thank the anonymous reviewers for their valuable suggestions. This work was supported by the BMBF-funded de.NBI Cloud within the German Network for Bioinformatics Infrastructure (de.NBI).

## Notes

### Competing Interest Statement

The authors have declared no competing interest.

