## Supplementary Material for "MOSGA 2: Comparative genomics and validation tools"

#### This PDF file includes:

Supplementary text
Tables S1 to S4 (not allowed for Brief Reports)
Figs. S1 to S6 (not allowed for Brief Reports)

### Supporting Information Text

#### EukCC

MOSGA 2 includes EukCC as a drop-in-replacement for the phylogenetic analysis and the genome completeness. Due to the underlying dependency as provided by the restricted licensed GeneMark, we put BUSCO into the foreground in our manuscript. EukCC will only appear in MOSGA 2 if the installing user provides the GeneMark binaries for the installation routine. However, EukCC can be used in the same way as BUSCO and generates as well a visualization about the genome completeness, which is shown in S1. We consider the best hit from the EukCC results for the visualization. As the BUSCO results in the main manuscript already indicate, the *Saccharomyces mikatae* genome seems to be incomplete.

#### Taxonomy search

Suppose the user prefers to choose MOSGA 2 for the closest and suitable configuration. In that case, MOSGA 2 includes a fast taxonomic search that identifies the most relative species from a given pool to a selected target species. The taxonomic search applies to every tool with pre-defined species-specific parameters. Our implementation builds on the NCBI taxonomy.

During the MOSGA 2 installation, we create an SQLite database with the scientific species names the corresponding NCBI taxonomy identifier. Additionally, we prepare the eukaryotic taxonomy tree with the taxonomy identifier (id) on trimmed NCBI taxonomy data, excluding all none-eukaryotic clades. In the MOSGA 2 GUI rules file, every tool with multiple prepared species-specific models contains taxonomy id. MOSGA 2 iterates through the tree for each search and counts the distance between each taxonomy id in the given pool of species and the target species. Initially, MOSGA 2 creates a short pathway list for all defined taxonomy ids from all tools to speed up the search. The species with the shortest distance are selected. MOSGA 2 weighs the lengths of taxonomic levels (kingdom, phylum, class, order, family, genus, and species) with a distance of 1 and clades to solve incomparable long branches inside the tree, with a distance value of 0.25.

As an example, we performed a taxonomy search for Augustus and BUSCO with *Taphrina alni* as the target species. An excerpt of the taxonomy search is shown in Figure S4 for Augustus. MOSGA 2 suggests using the *Pneumocystis jirovecii* model for *Taphrina alni* since these species have the closest distance. As the suitable BUSCO lineage, MOSGA 2 recommends using the Ascomycota, which represents the correct phylum for *Taphrina alni*.

#### Organellar DNA scanner

The organellar DNA scanner facilitates the identification of organellar scaffolds like mitochondria or plastids. It relatively ranks every scaffold to each other and sorts them according to the ranking. We tested this functionality by applying the scanner on twenty diverse eukaryotic genome assemblies and compared the suggested top scaffolds to the NCBI marked references. Table S4 shows the corresponding ranking position for each genome assembly classified by mitochondrial, plastids, and other organelles. An exemplary representation of a result table of the scanner is displayed in Table S3. To generate the plastid and mitochondrial reference protein databases, we clustered respective Refseqs databases with MMseqs2 SHA-0cc7e6 (`easy-cluster -s 7 -cluster-mode 1`) to identify core protein sets (minimum cluster size of 3000 for plastids and 50 for mitochondria). To reduce redundancy in these core gene sets, we performed an additional round of clustering with MMseqs2 at 80% identity (`easy-cluster -min-seq-id 0.8`) and only kept representative sequences of these clusters final database.

**Table S1. List of the used genome assemblies for the phylogenetic analysis with the corresponding NCBI assembly identifier and number of scaffolds.**

| Species | Genome Assembly | No. of scaffolds |
|---|---|---|
| *Saccharomyces arboricola* | GCA_000292725.1 | 35 |
| *Saccharomyces cerevisiae* | GCA_000146045.2 | 17 |
| *Saccharomyces eubayanusi* | GCA_001298625.1 | 24 |
| *Saccharomyces uvarum* | GCA_002242645.1 | 50 |
| *Saccharomyces paradoxus* | GCA_002079055.1 | 17 |
| *Saccharomyces kudriavzevii* | GCA_003327635.1 | 17 |
| *Saccharomyces mikatae* | GCA_000166975.1 | 1648 |

**Table S2. Strain and species name with the corresponding accession number of all used *Saccharomyces* strains for the protein-coding gene comparison.**

| Species and Strains | Genome Assembly |
| --- | --- |
| *Saccharomyces cerevisiae* HLJ167 | GCA_003271395.1 |
| *Saccharomyces cerevisiae* S288C | GCF_000146045.2 |
| *Saccharomyces cerevisiae* sake001 | GCA_001738375.1 |
| *Saccharomyces cerevisiae* SK1 | GCA_002057885.1 |
| *Saccharomyces cerevisiae* Y12 | GCA_015251845.1 |
| *Saccharomyces paradoxus* | GCA_002079055.1 |

**Table S3. Exemplary presentation of the organellar DNA scanner table results for *Nannochloropsis oceanica*. The number of partial and full ribosomal RNA, transfer RNA, matches of mitochondrial, and plastid genes, the percentage GC-content deviation as well the mitochondrial genes density and plastid genes density are taken into account for the scoring. GC-content deviations are only considered when they differ more than the standard deviation of the average GC-content. The first-ranked scaffold CP038136.1 represents the plastid genome and the second-ranked scaffold CP038136.1 the mitogenome.**

| Scaffold | rRNA | tRNA | rRNA(p) | Mito | Plastid | GC dev | GC | Len [kbp] | MD | PD | Score |
|---|---|---|---|---|---|---|---|---|---|---|---|
| CP038136.1 | 8 | 9 | 8 | 75 | 75 | 19.21* | 33.62 | 117.56 | 6.38 | 6.38 | 136838 |
| CP038137.1 | 1 | 5 | 1 | 25 | 25 | 20.97* | 31.87 | 38.07 | 6.57 | 6.57 | 51594 |
| CP038117.1 | 3 | 0 | 0 | 0 | 75 | 1.32 | 54.15 | 1596.16 | 0.0 | 0.47 | 36 |
| CP038119.1 | 0 | 1 | 0 | 0 | 50 | 1.25 | 54.08 | 723.33 | 0.0 | 0.69 | 35 |
| CP038107.1 | 1 | 0 | 0 | 0 | 54 | 1.28 | 54.11 | 1159.84 | 0.0 | 0.47 | 25 |
| CP038130.1 | 1 | 3 | 0 | 0 | 41 | 1.11 | 53.94 | 1534.45 | 0.0 | 0.27 | 12 |
| CP038122.1 | 0 | 0 | 0 | 0 | 25 | 1.84 | 54.67 | 608.09 | 0.0 | 0.41 | 10 |
| CP038115.1 | 0 | 2 | 0 | 0 | 25 | 1.0 | 53.83 | 837.05 | 0.0 | 0.3 | 8 |
| CP038111.1 | 7 | 0 | 0 | 0 | 25 | 1.7 | 54.53 | 961.38 | 0.0 | 0.26 | 8 |
| CP038118.1 | 0 | 0 | 0 | 0 | 25 | 0.7 | 53.54 | 814.1 | 0.0 | 0.31 | 7 |
| CP038106.1 | 0 | 0 | 0 | 25 | 44 | 1.19 | 54.03 | 1670.64 | 0.15 | 0.26 | 2 |
| CP038132.1 | 3 | 1 | 0 | 25 | 18 | 0.49 | 53.32 | 1340.16 | 0.19 | 0.13 | 1 |
| CP038128.1 | 0 | 1 | 0 | 27 | 30 | 0.98 | 53.81 | 1579.29 | 0.17 | 0.19 | 1 |
| CP038126.1 | 0 | 1 | 0 | 0 | 0 | 1.33 | 54.16 | 475.66 | 0.0 | 0.0 | 1 |
| CP038124.1 | 0 | 0 | 0 | 0 | 10 | 1.62 | 54.45 | 551.95 | 0.0 | 0.18 | 1 |
| CP038120.1 | 1 | 0 | 0 | 0 | 0 | 1.63 | 54.46 | 645.58 | 0.0 | 0.0 | 1 |
| CP038109.1 | 1 | 0 | 0 | 14 | 28 | 1.62 | 54.45 | 1145.27 | 0.12 | 0.24 | 1 |

**Table S4. Summarized results from the organellar DNA scanner of twenty eukaryotic genome assemblies. The positions of each verified organellar scaffold from single organellar DNA scanner results are shown. Empty positions are resulting from the absence of known organellar scaffolds inside an assembly. Asterisks mark problematic assemblies that contain multiple unplaced scaffolds.**

| Species | Assembly | Unplaced Contigs | No. Scaffolds | Score position | | |
|---|---|---|---|---|---|---|
| | | | | Mitochondrium | Chloroplast | Other organelles |
| *Arabidopsis thaliana* | GCA_000001735.2 | No | 7 | 1 | 2 | |
| *Apis mellifera* | GCA_003254395.2 | Yes | 11 | 1 | | |
| *Babesia microti* | GCA_000691945.2 | No | 6 | 2 | | 1 |
| *Bos taurus* | GCA_002263795.2 | Yes | 2211 | 1 | | |
| *Caenorhabditis elegans* | GCA_000002985.3 | No | 7 | 1 | | |
| *Cafeteria burkhardae* | GCA_008330645.1 | Yes | 170 | 1 | | |
| *Cardiosporidium cionae* | GCA_015476325.1 | Yes | 2204 | | | 3 |
| *Corvus cornix* | GCA_000738735.2 | Yes | 113 | 1 | | |
| *Danio rerio* | GCA_000002035.4 | Yes | 1917 | 1 | | |
| *Drosophila melanogaster* | GCA_000001215.4 | Yes | 1870 | 1 | | |
| *Homo sapiens* | GCA_000306695.2 | No | 164 | 1 | | |
| *Ipomoea triloba* | GCA_003576645.1 | No | 17 | | 1 | |
| *Nannochloropsis oceanica* | GCA_004519485.1 | No | 32 | 1 | 2 | |
| *Plasmodium falciparum* | GCA_000002765.3 | No | 15 | | | 1 |
| *Prunus dulcis* | GCA_902201215.1 | Yes | 692 | | 2 | |
| *Saccharomyces cerevisiae* | GCA_000146045.2 | No | 17 | 1 | | |
| *Salmo salar* | GCA_000233375.4 | Yes | 241573 | 6 | | |
| *Strongylocentrotus purpuratus* | GCA_000002235.4 | Yes | 871 | 1 | | |
| *Tribolium castaneum* | GCA_000002335.3 | Yes | 2149 | 7 | | |
| *Zea mays* | GCA_902167145.1 | Yes | 687 | 27 | | 5 |

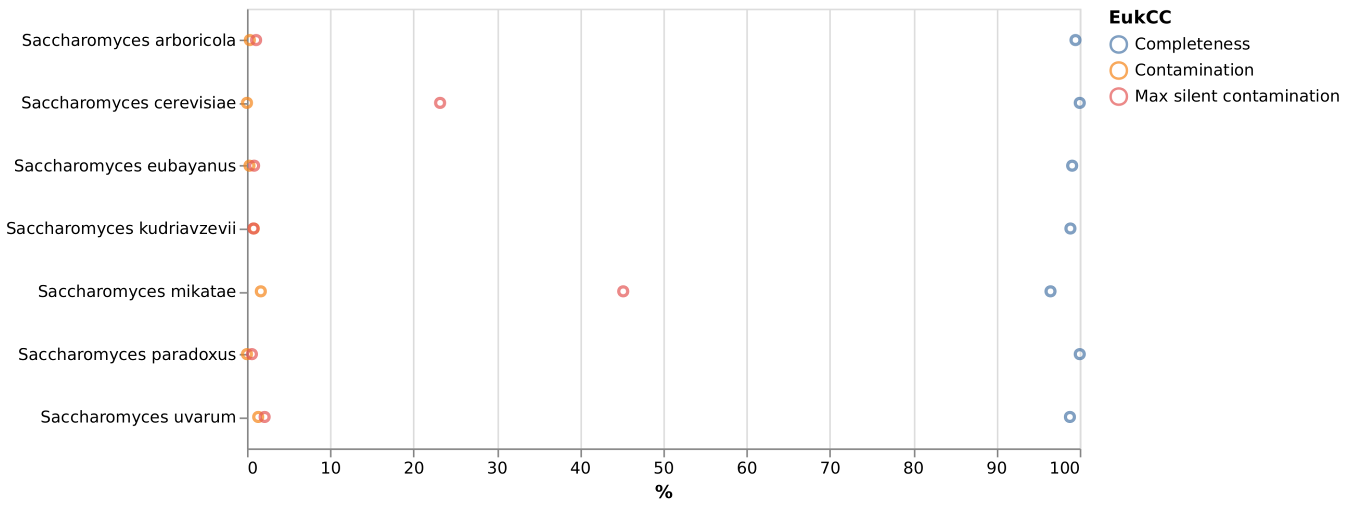


**Fig. S1.** EukCC analysis. Summarized results of a EukCC analysis on seven yeast species show that all genome assemblies are mostly complete. *S. cerevisiae* and *S. mikatae* contain putative high silent contamination that matches for *S. mikatae* analysis results from BUSCO.

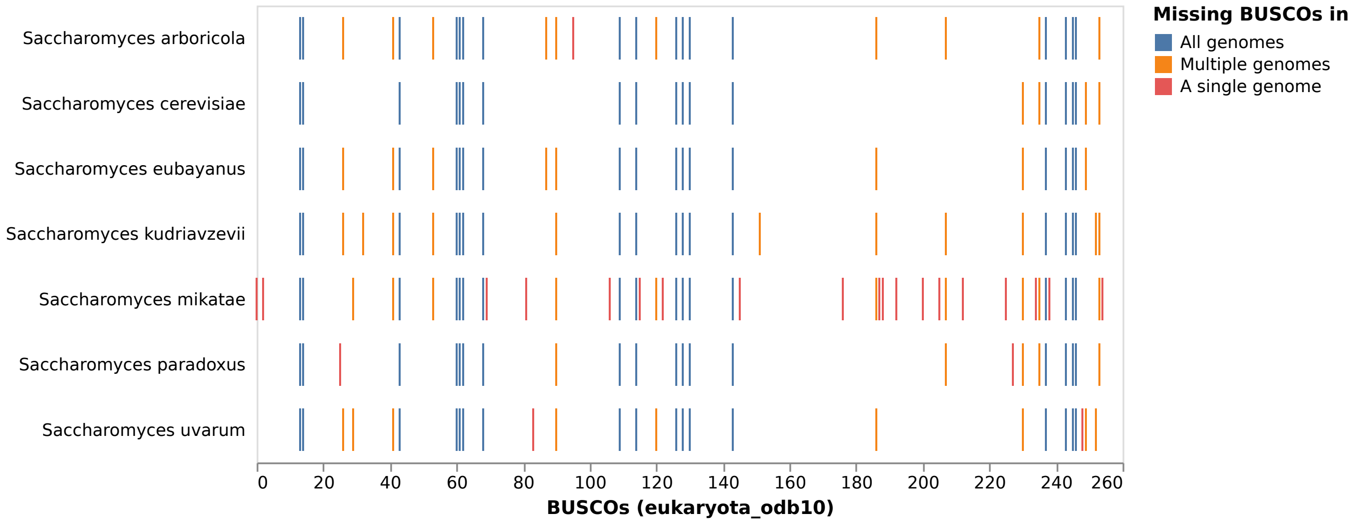


**Fig. S2.** Missing BUSCOS. The stripe chart represents the uniqueness and the commonality of missing BUSCOs. All genomes order the Eukaryota lineage BUSCO equally.

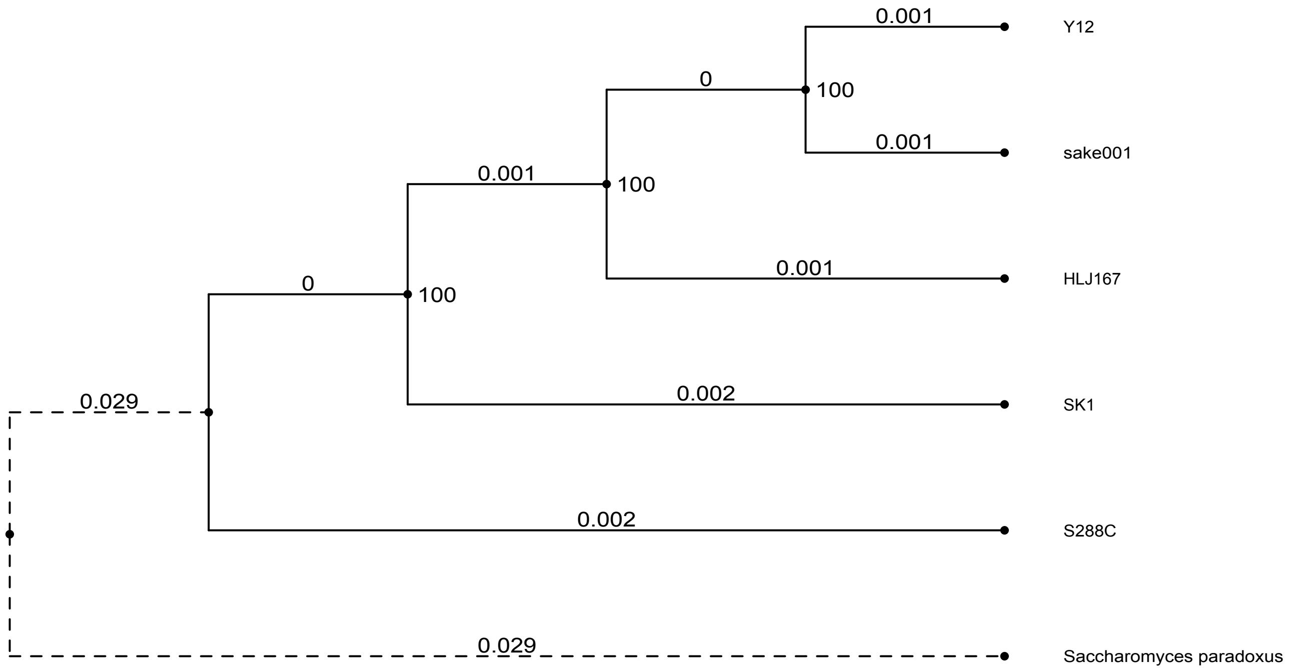


**Fig. S3.** Phylogenetic tree analysis on six different *Saccharomyces* strains and species. This phylogenetic tree analysis is based on 1837 common BUSCO genes. MAFFT performs the Multiple sequence alignment and traimAl the trimming. RAxML computes with phylogenetics in the GTRCAT mode and calculates the support values through bootstrapping.

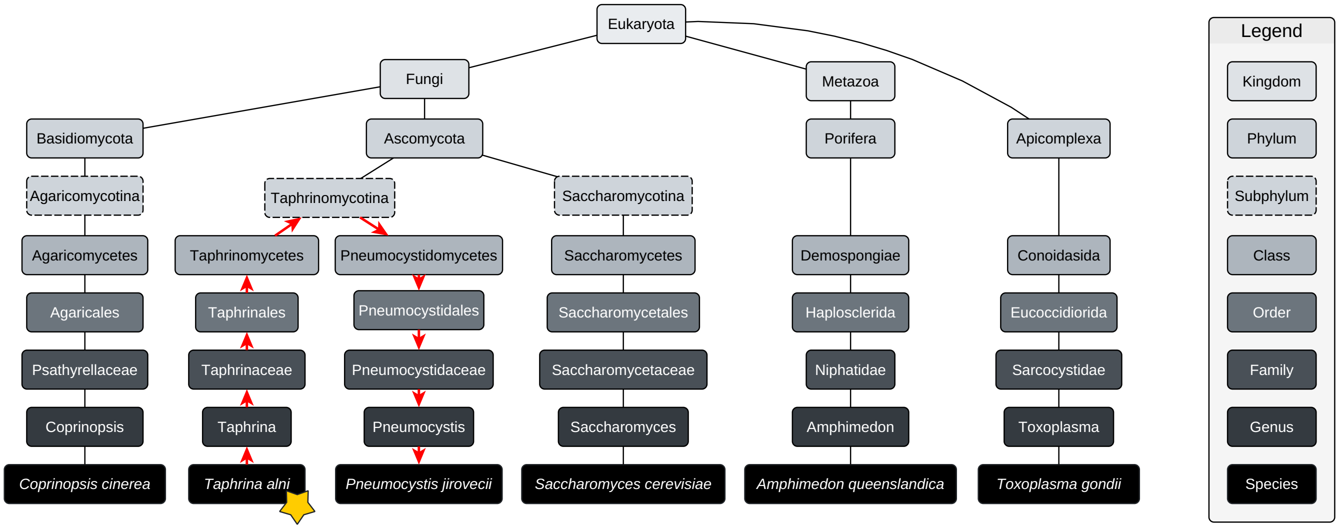


**Fig. S4.** An excerpt of a visual taxonomy tree representation for the taxonomic search. MOSGA 2 builds the whole taxonomy tree for the Eukaryota superkingdom and searches the target species inside this tree. In this example, *Taphrina alni* represents the target species marked with an asterisk. MOSGA 2 computes every distance to each available taxon in the models (Augustus) or lineage (BUSCO) list and selects the shortest distance in the following step, in this case, *Pneumocystis jirovecii*. Red arrows highlight the chosen path. We do not show the clades in between the taxonomical, which were also considered and less weighted.

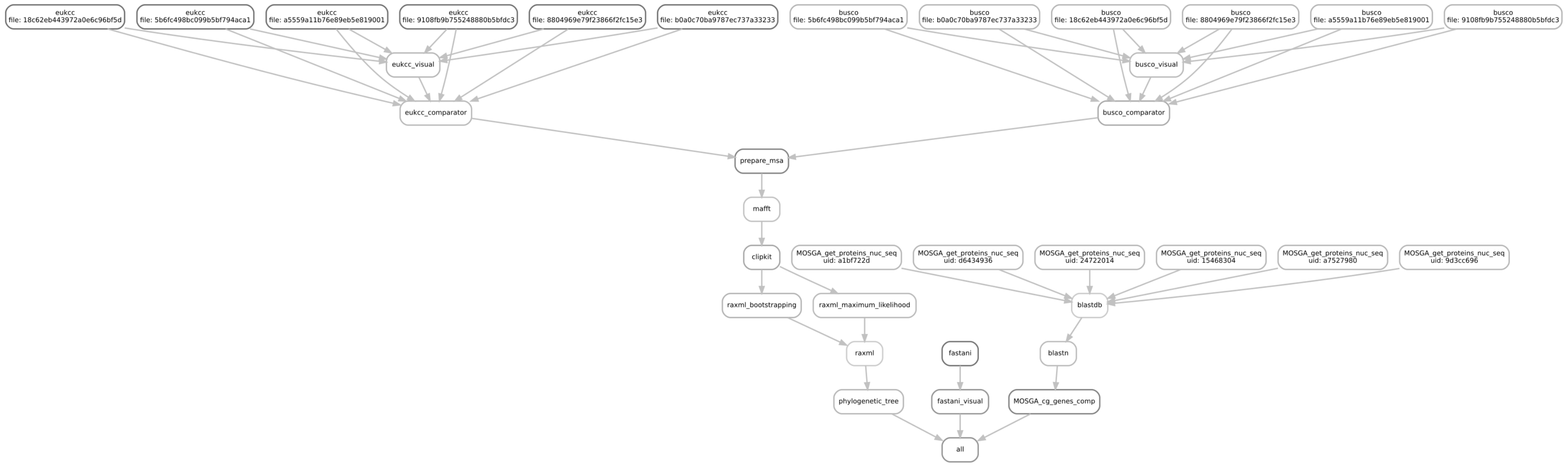


**Fig. S5.** Exemplary pipeline run for the comparative genomics workflow, including BUSCO, EukCC, MAFFT, ClipKit, RAxML, and the protein-coding gene comparison.

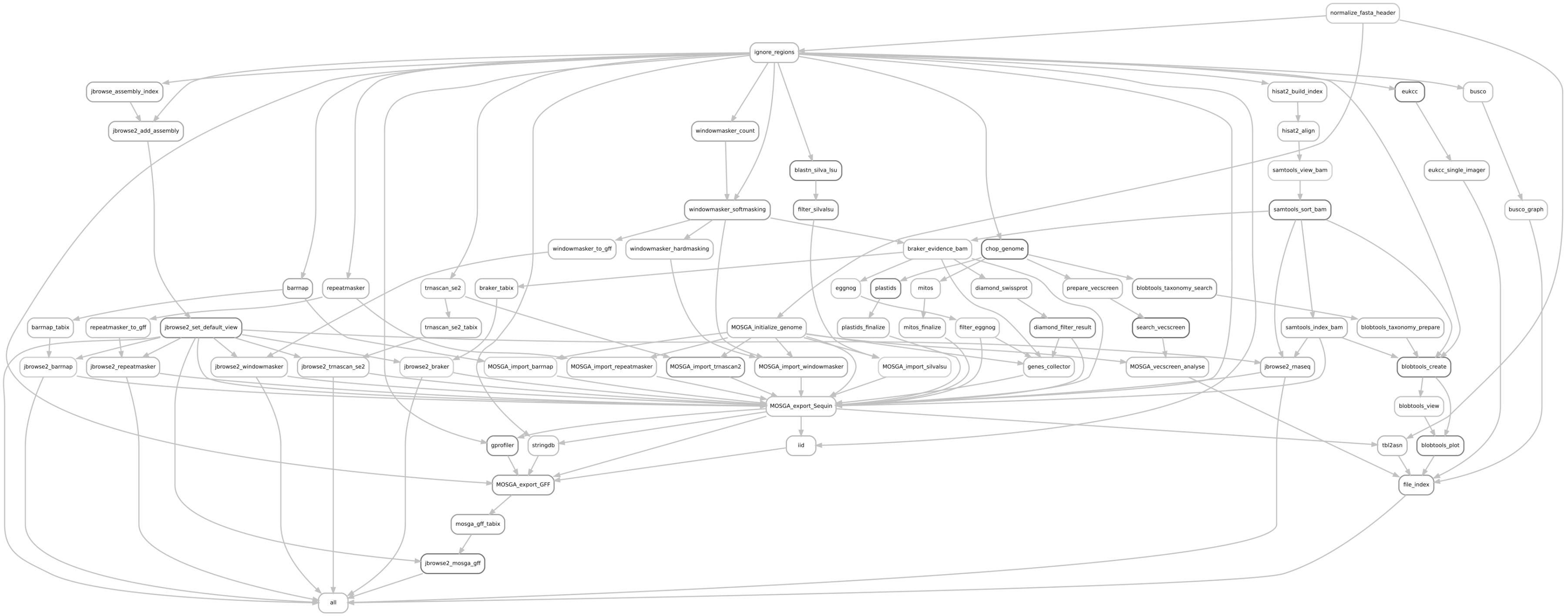


**Fig. S6.** Exemplary pipeline run for the annotation workflow, including RNA-seq preprocessing by HiSat2, Samtools, BRAKER2, tRNAscan-SE2.0, SILVIA, barrnap, RepeatMasker, Windowmasker, JBrowse, BUSCO, EukCC, BlobTools, VecScreen, tbl2asn, the organellar DNA scanner, functional annotation with EggNog and Swiss-Prot and various scripts, external APIs and visualization creation.
